# The separation of the effects of atomic displacement and local resolution in crystallographic and cryo-electron microscopy maps

**DOI:** 10.1101/2025.07.21.665840

**Authors:** Vladimir Y. Lunin, Natalia L. Lunina

## Abstract

Uncertainties in atomic coordinates and loss of resolution blur atomic images on density maps. The resolution cutoff leads to an additional effect, the appearance of Fourier ripples. Ignoring these ripples makes it impossible to distinguish between the two sources of image blurring. However, separate determination of the atomic displacement parameter and local resolution becomes possible if an advanced method for calculating the image of the atom in the observed map is used, which includes modeling of ripples.

**Synopsis:** Shell-based density modelling allows us to separate the effects of atomic displacement and of resolution cutoffs in crystallographic and cryo-electron microscopy maps during refining atomic models in real space.

## 1. Introduction

In what follows, the density distribution of the object under study refers to a three-dimensional electron density distribution in X-ray crystallography (XR) or to the distribution of electrostatic scattering potential in cryo-electron microscopy (cryo-EM), or to any other similar spatial field. Let *ρ*^*obs*^(**r**) be a map of this distribution obtained from the experiment. This map can be calculated as a Fourier series in XR, or as an image reconstruction in cryo-EM. These maps are affected by various uncertainties, such as dynamic and static uncertainties in the positions of atomic centers, reduced resolution, varying across the map, sample instability, arbitrary scaling, background, and so on.

Let *ρ*^*theor*^(**r**) be the corresponding theoretical density distribution calculated from an atomic model of the respective structure, ignoring both atomic uncertainties and loss of resolution. To compare, point-by-point, the map calculated from the atomic model with the observed one, the former must reproduce the distortions present in the latter. We denote by *ρ*^*calc*^(**r**) such a map obtained from *ρ*^*theor*^(**r**) after introducing the respective distortions. Real-space refinement, first introduced by Diamond (1971), searches for the parameters of both the model and the distortion by minimizing the discrepancy between *ρ*^*calc*^(**r**) and *ρ*^*obs*^(**r**).

The concept of ‘resolution’ characterizes the details seen in a map (e.g., Urzhumtseva et al., 2013), and the respective value may vary from one map region to another, thus, a single value for the entire map may be insufficient (Cardone *et al*., 2013). When calculating a map as the sum of atomic images, the local resolution can be associated with each atom. (Urzhumtsev & Lunin 2022). Here we define the local resolution of an atomic image as the value of the cutoff applied to the inverse Fourier transform when calculating the image from the atomic scattering factor. Section 2 precises this definition.

## 2. The image of an atom in the observed map

In XR and cryo-EM, the theoretical density distribution is usually assumed to be a sum of independent atomic contributions (e.g., Urzhumtsev & Lunin, 2019)

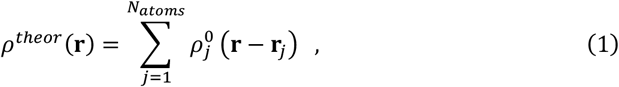

where *N*_*atoms*_ is the number of atoms in the object under study, **r**_*j*_ are the coordinates of the atomic centers, and 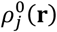 is the theoretical density distribution of atom *j* placed at the origin. The theoretical density distribution is usually approximated by a sum

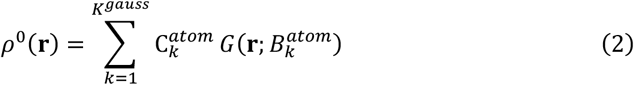

of a few normalized Gaussian function in three-dimensional space **R**^3^

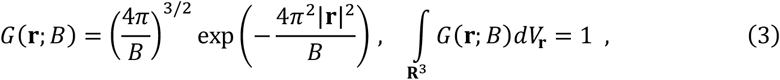

with known coefficients 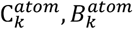 (e.g., Doyle & Turner, 1968; Peng, 1999; Brown et al., 2006). Here, and in what follows, to simplify the formulas, we consider *K*^*gauss*^ = 1, *i*.*e*., the so-called ‘Gaussian atom’. The results obtained are directly generalized to the case of a multi-Gaussian theoretical density (2).

### 2.1. Distortion due to the positional uncertainty of an atom

In both XR and cryo-EM, the experimental map is obtained by averaging over a huge number of copies of the object under study. These copies are almost identical, but they may have equivalent atoms in slightly different positions. When for some atom, the uncertainty of its position is described by a probability distribution *P*(**r**), then its expected image, smeared by this uncertainty, is given by convolution.

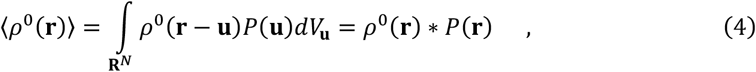

Here and elsewhere the symbol * denotes the mathematical operation of convolution. When working with medium-resolution maps, this probability distribution is usually given by an isotropic Gaussian function *P*(**r**) = *G*(**r**; *B*) with a parameter *B* called the Atomic Displacement Parameter (ADP). The image of the atom, distorted by positional uncertainty, takes the form of

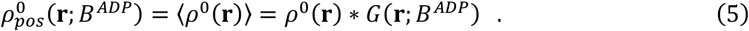

### 2.2. Distortion due to local loss of resolution

The scattering factor of an atom is the three-dimensional Fourier transform of its atomic density distribution

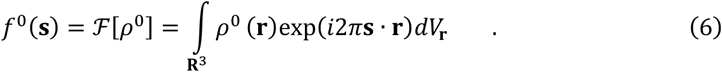

The density distribution can be reconstructed from the scattering factor using the inverse Fourier transform

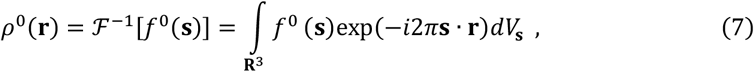

When the last integral is calculated over of a sphere of radius *s*_*max*_ = *D*^−1^

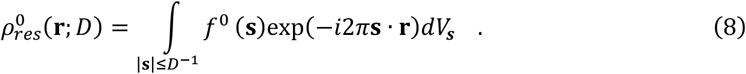

we call the result ‘the image of an atom at the resolution D’. This image can be equivalently represented by a convolution

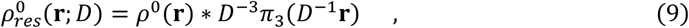

where *π*_3_(**r**) is the normalized three-dimensional interference function

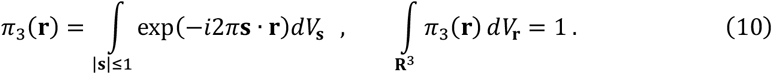

The explicit form of this function is

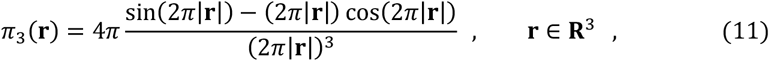

(Fig. 1). For a point atom (an atom with a density distribution described by the Dirac delta function), its image at the resolution D is a scaled interference function

**Fig. 1.**
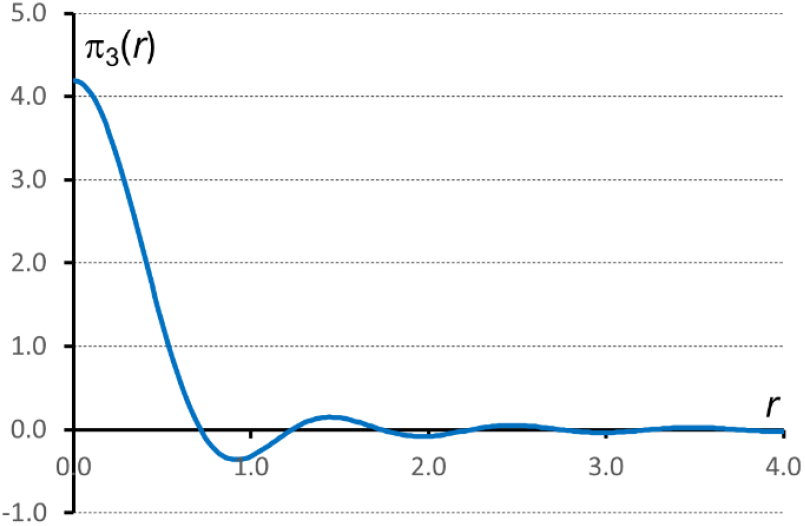
The normalized 3D interference function.

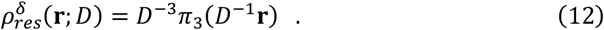

The effect of the resolution cut-off results in both blurring the central peak and appearance of spherical shells with alternating positive and negative function values.

The two types of principal distortions of the theoretical density, (5) and (9), can be combined into a single expression

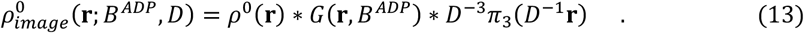

### 2.3. Gaussian real-space refinement

The relatively small amplitude of the ripples in the interference function suggests the idea of approximating the interference function by a Gaussian function, ignoring the ripples (e.g., Mooij et al., 2006; Sorzano et al., 2015)

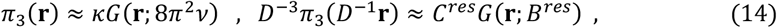

Where

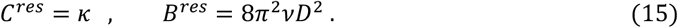

For a Gaussian atom, its image (13) is a convolution of Gaussian functions. Due to the convolution property of Gaussian functions, the result is also a Gaussian function

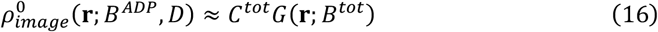

With

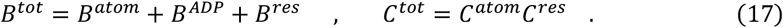

When using Gaussian models for atom density, atomic displacement, and loss of resolution, the calculated density takes the form

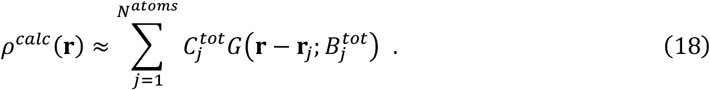

An attempt to fit this function to *ρ*^*obs*^(**r**) by real-space refinement allows us to find for each atom, beside the coordinates, only the cumulative values 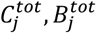 of distortion parameters, but not the separate values 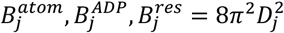, linked by (17).

## 3. Shell functions

Recently, a method for efficient calculation of the convolution (13) has been proposed (Urzhumtsev & Lunin, 2022; Urzhumtseva et al., 2023; Urzhumtsev & Lunin, 2024). This method is based on an approximation to the normalized interference function by the sum of shell functions

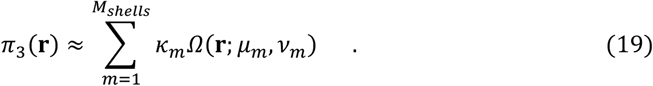

The shell function *Ω*(**r**; *μ, ν*) is defined as the convolution of a singular uniform density distribution on the spherical surface with a Gaussian distribution. This function depends on two parameters: the radius *μ* of the spherical shell, around which the density is concentrated, and a parameter *ν* that regulates the spread of the function around the shell. In three-dimensional space, the shell function can be written as

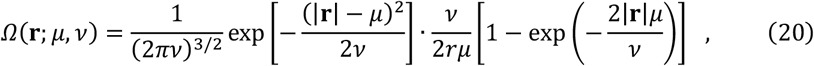

and possesses several properties including

- a Gaussian function is a special case of a shell function

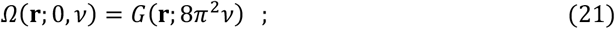
- changing the scale of the argument can be reduced to a modification of its parameters

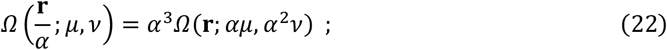
- the convolution of a shell function with a Gaussian function again results in a shell function with modified parameter values.

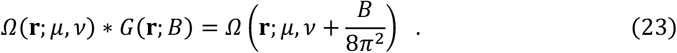

Due to the convolution property (23), the image of a Gaussian atom *ρ*^0^(**r**) = *C*^*atom*^*G*(**r**, *B*^*atom*^) distorted by the positional uncertainty *B*^*ADP*^ and the resolution *D* has a form

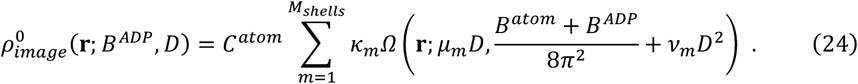

Reducing the approximation (19) only to the first term, for which *μ*_1_ = 0, we obtain a Gaussian approximation (14-15) with the constants *ν*_1_ = 0.149577, κ_1_ = 3.974732 (Urzhumtsev & Lunin, 2024). In other words, additional blurring of the central peak of an image of a Gaussian atom due to resolution cutoff may be modeled by increasing the ADP by

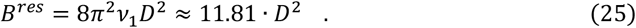

Adding the second term from (a20) allows us to take into account the first negative ripple

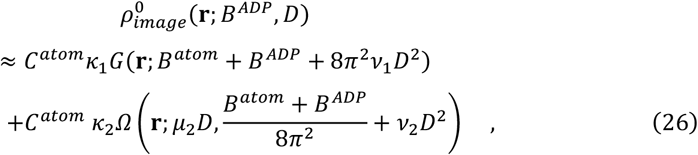

where *μ*_2_ = 0.839261, *ν*_2_ = 0.081904, κ_2_ = −4.722464.

## 4. Shell-based refinement

Now let the experimental map be a sum of atomic contributions where each of them is a sum of two terms, a Gaussian one with *μ*_1_ = 0, and a shell function with *μ*_2_ ≠ 0:

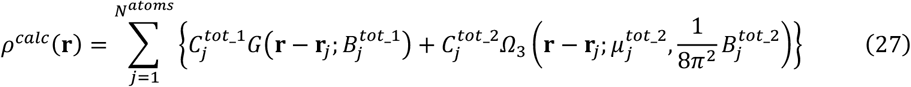

This time, searching for the best fit of the model map *ρ*^*calc*^(**r**) to *ρ*^*obs*^(**r**), as above, one can find, in particular, the values of 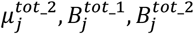 such, that

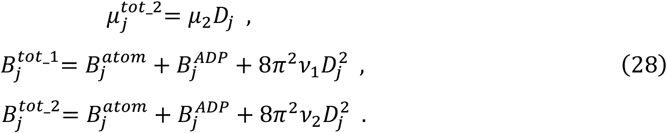

Now, with three found values, we can determine the value of *D*_*j*_ in two ways:

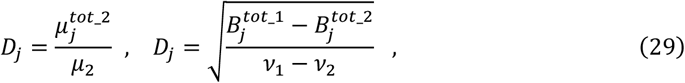

where *ν*_1_, *ν*_2_, *μ*_2_ are the fixed values of the parameters of expansion (19). The found resolution values *D*_*j*_ make it possible in turn to remove the components 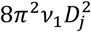 and 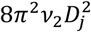 from equations (28) and, with known atomic scattering factors, determine the values of the individual parameters 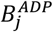.

## 5. Conclusions

The performed analysis shows that real-space refinement allows us to find separately displacement and local resolution parameters, if at least one shell function is added to an approximation of atomic images. Intuitively, increasing the number of terms in (24) will provide more accurate information about the parameters, refining their values. The possibility of independently determining parameters *B*^*ADP*^ and *D* when using the shell approximation (24) of the atomic image has been demonstrated through a series of tests by Lunin et al. (2023). Fig. 2 illustrates shell-based determination of atomic displacement parameters and local resolution for the nucleosome-sirtuin complex.

**Fig. 2.**
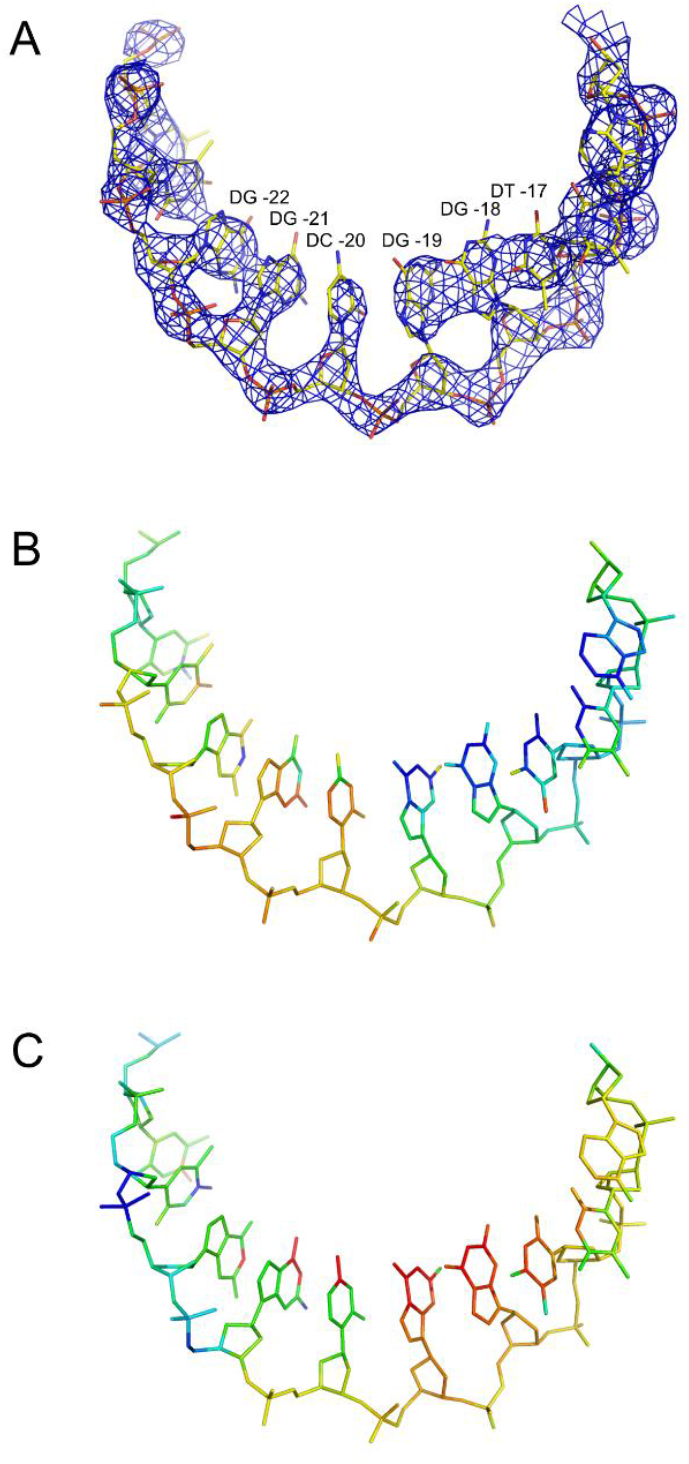
A fragment of the DNA chain J in the nucleosome-sirtuin complex (Smirnova et al., 2023). A. The atomic model (PDB entry code 8of4) is superimposed on the cryo-EM map (emd_16845.map) B. The atomic model colored according to the *B*^*ADP*^ values found with the shell-based density model: blue for smaller values starting at 0. Å^2^, red for higher values up to 100. Å^2^. C. The atomic model colored according to the local resolution values *D* found: blue indicates lower D-values (higher resolution, starting at 2.5 Å); red indicates higher D values (lower resolution, up until 4.5 Å). Figures were prepared using *PyMOL* software (Schrödinger & De Lano, 2020).

The authors thank A.G. Urzhumtsev for valuable discussions.

## References

Brown, P. J., Fox, A. G., Maslen, E. N., O’Keefe, M. A. & Willis, B. T.M. (2006). International Tables for Crystallography (2006). Vol. C, Chapter 6.1, pp. 554–595.

Cardone, G., Heymann, J. B. & Steven, A. C. (2013). J. Struct. Biol. 184, 226–236. 10.1016/j.jsb.2013.08.002

Diamond, R. (1971). Acta Cryst. A27, 436–452. 10.1107/S0567739471000986

Doyle, P. A. & Turner, P. S. (1968). Acta Cryst. A 24, 390–397. 10.1107/S0567739468000756

Lunin V.Y., Lunina N.L., Urzhumtsev A.G. (2023). Current Research in Structural Biology, 6, 100102. 10.1016/j.crstbi.2023.100102

Mooij, W. T. M., Hartshorn, M. J., Tickle, I. J., Sharff, A. J., Verdonk, M. L. & Jhoti, H. (2006). Chem. Med. Chem, 1, 827–838. 10.1002/cmdc.200600074

Peng, L.-M. (1999). Micron. 30, 625–648. 10.1016/S0968-4328(99)00033-5

Schrödinger, L. & DeLano, W. (2020). PyMOL, Available at: http://www.pymol.org/pymol.

Smirnova, E., Bignon, E., Schultz, P., Papai, G. & Ben-Shem, A. (2023). eLife, 10.7554/eLife.87989.3

Sorzano, C. O. S., Vargas, J., Otón, J., Abrishami, V., de la Rosa-Trevín, J. M., del Riego, S., Fernández-Alderete, A., Martínez-Rey, C., Marabini, R. & Carazo, J. M. (2015). AIMS Biophys. 2, 8–20. 10.3934/biophy.2015.1.8

Urzhumtsev, A. & Lunin, V.Y. (2022). IUCrJ. 9, 728–734 10.1107/S2052252522008260

Urzhumtsev, A.G. & Lunin V.Y. (2024). 2412.14350v1. 10.48550/arXiv.2412.14350

Urzhumtseva, L., Klaholz, B. & Urzhumtsev, A. (2013). Acta Cryst. D69, 1921–1934. 10.1107/S0907444913016673

Urzhumtseva, L., Lunin, V.Y. & Urzhumtsev, A.G. (2023). J. Appl. Cryst. 56, 302–311. 10.1107/S160057672201144X

